# Enhancing Rice Growth and Yield with Weed Endophytic Bacteria *Alcaligenes faecalis* and *Metabacillus indicus* Under Reduced Chemical Fertilization

**DOI:** 10.1101/2023.12.18.572215

**Authors:** Kaniz Fatema, Nur Uddin Mahmud, Dipali Rani Gupta, Md. Nurealam Siddiqui, Tahsin Islam Sakif, Aniruddha Sarker, Andrew G Sharpe, Tofazzal Islam

## Abstract

Endophytic bacteria, recognized as eco-friendly biofertilizers, have demonstrated the potential to enhance crop growth and yield. While the plant growth-promoting effects of endophytic bacteria have been extensively studied, the impact of weed endophytes remains less explored. In this study, we aimed to isolate endophytic bacteria from native weeds and assess their plant growth-promoting abilities in rice under varying chemical fertilization. The evaluation encompassed measurements of mineral phosphate and potash solubilization, as well as indole-3-acetic acid (IAA) production activity by the selected isolates. Two promising strains, tentatively identified as *Alcaligenes faecalis* (BTCP01) from Eleusine indica (Goose grass) and *Metabacillus indicus* (BTDR03) from *Cynodon dactylon* (Bermuda grass) based on 16S rRNA gene phylogeny, exhibited noteworthy phosphate and potassium solubilization activity, respectively. BTCP01 demonstrated superior phosphate solubilizing activity, while BTDR03 exhibited the highest potassium (K) solubilizing activity. Both isolates synthesized IAA in the presence of L-tryptophan, with the detection of *nifH* and *ipdC* genes in their genomes. Application of isolates BTCP01 and BTDR03 through root dipping and spraying at the flowering stage significantly enhanced the agronomic performance of rice variety BRRI dhan29. Notably, combining both strains with 50% of recommended N, P, and K fertilizer doses led to a substantial increase in rice grain yields compared to control plants receiving 100% of recommended doses. Taken together, our results indicate that weed endophytic bacterial strains hold promise as biofertilizers, potentially reducing the dependency on chemical fertilizers by up to 50%, thereby fostering sustainable rice production.

## 1. Introduction

Rice (*Oryza sativa* L.), a staple for nearly half the global population and the third-largest cereal crop worldwide, holds paramount importance in sustaining human diets [1]. In 2020, the USDA reported global rice production at 503.17 million metric tons, utilizing 11% of cropland [2]. Bangladesh, ranking third in global rice production, dedicates around 78% of its arable land to rice cultivation, projecting an output of 38.4 million tons [3]. For developing countries, including Bangladesh, rice contributes significantly to daily caloric intake, providing 27% of dietary energy, 20% of dietary protein, and 3% of dietary fat [4]. However, the heavy reliance on agrochemicals, such as urea, triple super phosphate (TSP), and muriate of potash (MoP) for the higher yield of rice, poses environmental threats, prompting the exploration of sustainable alternatives [6]. Furthermore, the natural mineral sources for the production of these three major fertilizers required for rice production are finite and depleting day by day.

This study addresses the urgent need for low-cost technologies to enhance crop productivity while mitigating the environmental impact of chemical fertilizers. While various strategies exist, leveraging beneficial microbes offers an economical and viable solution [10,11,12]. Notably, plant growth-promoting bacteria (PGPB) or plant probiotic bacteria have emerged as promising contributors to enhanced productivity, particularly in rice cultivation [13]. A large body of literature indicate that plant-associated bacteria as biofertilizers and/biostimulants are natural and renewable bioresources for the reduction of hazardous synthetic chemicals required rice production. The mechanisms of the plant probiotic bacteria include fixation of atmospheric nitrogen, solubilization of plant essential nutrient elements in soils, production of phytohormones and various metabolites and regulation of gene expression in the host plants. Some of these plant probiotic bacteria belonging to the genera of *Bacillus*, *Rhizobium*, *Pseudomonas*, *Enterobacter*, *Paraburkholderia, Delftia* etc. have been proven as biofertilizers and/or biopesticides in production of many crops including rice [10-21].

Despite the valuable role of plant endophytic bacteria in plant growth, their potential, especially weed endophytes, remains poorly underexplored [13-33]. The hypothesis of this study was weed endophytes from rice field can enhance growth and yield of rice under low fertilization conditions. This research aims to isolate and characterize endophytic bacteria from rice-associated weeds, assess their impact on rice growth and yield, identify the bacteria through 16S rRNA gene sequencing, and elucidate their growth-promoting roles by detecting genes involved in nitrogen fixation, phosphorus and potassium solubilization, and IAA production. While weed endophytes are often overlooked, their adaptation to diverse conditions makes them potential reservoirs of beneficial bacteria with unique capabilities, offering novel insights for sustainable agriculture.

## 2. Materials and Methods

### 2.1. Experimental sites

The native weed samples were collected to isolate bacteria from the field laboratory at Bangabandhu Sheikh Mujibur Rahman Agricultural University (BSMRAU), located in Gazipur, Bangladesh (24.09° N and 90.25° E).

### 2.2. Collection of plant materials and isolation of bacteria

Root and shoot samples were collected from various weed species, including Ulu (Cogon grass: *Imperata cylindrica*), Chapra (Goose grass: *Eleusine indica*), Kashful (wild sugarcane: *Saccharum spontaneum*), Mutha (Nutsedge: *Cyperus rotundus*), Durba (Bermuda grass: *Cynodon dactylon*), Anguli Ghash (Scrab grass: *Digitaria sanguinalis*), Khude sama (Jungle Rice: *Echinochloa colonum*), Arail (Swamp rice grass: *Leersia hexanda* Sw.), Boro sama (Burnyard Grass: *Echinochloa crussgalli*), Kakpaya (Crow foot grass: *Dactyloctenium aegyptium*). These samples were collected at the vegetative stage from experimental sites where the weeds naturally grew, facilitating bacterial isolation. Additionally, seeds of the rice variety CV. BRRI dhan29 were procured from the Bangladesh Rice Research Institute (BRRI) for use in pot experiments to isolate endophytic bacteria. To prepare the root and shoot samples for isolation, thorough washing with distilled water, followed by a 5-minute rinse with 70% ethanol, was conducted. Subsequently, the samples were sterilized with 1% NaOCl for 1 minute, followed by a thorough rinse with sterile distilled water. The tissue was then further rinsed for 1 minute in 100% ethanol, followed by five washes with sterile distilled water. The processed samples were crushed in a sterilized mortar and pestle, diluted with sterile distilled water (SDW) up to a 1 × 10^-6^ dilution. A 100 µl aliquot of each dilution was evenly spread on Petri dishes containing nutrient broth agar medium (NBA) and incubated for 2 days at 25°C [10]. Colonies with distinct appearances were then transferred to new nutrient broth agar medium plates for purification. The purified isolates (single colonies) were preserved in a 20% glycerol solution at −20°C.

### 2.3. Seedling assay

The bacterial strains were initially cultured in 250 mL conical flasks containing 200 mL of NB (Nutrient Broth) medium, placed on an orbital shaker at 120 rpm, and incubated for 72 hours at 27°C. Subsequently, the resulting broth underwent centrifugation at 15,000 rpm for 1 minute at 4°C, and the bacterial cells were collected and washed twice with sterilized distilled water (SDW). The bacterial pellets were then resuspended in 0.6 mL of SDW, vortexed for 45 seconds, and prepared for seed treatment.

For seed treatment, 1 gram of surface-sterilized rice seeds (cv. BRRI dhan29) was immersed in the bacterial suspension, dried overnight at room temperature, and arranged on a Petri dish with water-soaked sterilized filter paper. Following seed germination, the seedlings were allowed to grow for two weeks, receiving alternate-day watering. Germination percentages were calculated at two days after inoculation (DAI). After 15 DAI, the impact of plant probiotic bacteria on rice seedling growth was evaluated, recording parameters such as germination rate, shoot length (cm), root length (cm), shoot fresh weight (g), and root fresh weight (g).

### 2.4. Biochemical characterization of isolated bacteria

For the biochemical characterization of the isolated bacteria, the gram reaction was determined according to the method outlined by [35]. Various biochemical tests were conducted to characterize the isolated bacteria, following the criteria outlined by Bergey et al. (1994). The assessment of KOH solubility involved mixing bacterial isolates with a 3% KOH solution on a clean slide for 1 minute, and the observation of a thread-like mass. Catalase and oxidase tests were performed following the procedures described by [36,37].

### 2.5. DNA extraction, 16S rRNA gene amplification and phylogenetic analysis of isolated bacteria

The bacterial DNA extraction utilized the lysozyme-SDS-phenol-chloroform method with phenol-chloroform-isoamyl alcohol (25:24:1), followed by precipitation with isopropanol, following the procedure outlined by Maniatis et al. (1982). Subsequently, the extracted DNA underwent treatment with DNase-free RNase (Sigma Chemical Co., St. Louis, MO, USA) at a final concentration of 0.2 mg/ml, incubated at 37°C for 15 minutes. Amplification of the 16S rRNA gene was achieved using a universal primer (27F, 5’AGAGTTTGATCCTGGCTCAG3’; 1492R, 5’GGTTACCTGTTACGACTT3’) (Reysenbach et al., 1992), and the reaction was carried out in a thermocycler (Mastercycler® Gradient, Eppendorf, Hamburg, Germany) following established guidelines.

The amplified products underwent purification using Quick PCR purification columns (Promega, Madison, WI, USA) and were subsequently sequenced with the Big Dye Terminator Cycle Sequencing Ready Reaction Kit on an Applied Biosystems analyzer (Applied Biosystems, Forster City, CA, USA). Sequences were compared to the NCBI GenBank database (http://www.ncbi.nlm.nih.gov) through a BLASTN search. For phylogenetic analysis, reference sequences were retrieved, and multiple sequence alignment was conducted using the CLUSTALW program in BioEdit version 7.2.3 [38], with manual editing of gaps. The construction of a phylogenetic tree employed the neighbor-joining method (NJ) (Saitou and Nei, 1987) in the MEGA software package version MEGA7 (Kumar et al., 2016). Pair-wise evolutionary distances were calculated using the Maximum Composite Likelihood method [39], and confidence values, based on sequence grouping, were obtained through bootstrap analysis with 1000 replicates [40].

### 2.6. Design of primers for amplification of nifH and ipdC genes

Primers were meticulously crafted through homology searches for a specific gene *(nifH* and *ipdC*) within *Alkaligenes* spp. and *Metabacillus* spp., as documented in the NCBI GenBank. The primer pairs were designed based on the region exhibiting homology across these genera (refer to Table S1).

### 2.7. Bioassays for plant growth promoting traits

#### 2.7.1. Determination of IAA production

The determination of indole-3-acetic acid (IAA) production by two bacterial isolates followed the original protocol proposed by [41], with minor adaptations. In brief, isolated colonies were inoculated into 50 ml of sterile Jensen broth (comprising 20 g/l sucrose, 1 g/l K2HPO4, 0.5 g/l MgSO4 • 7H2O, 0.5 g/l NaCl, 0.1 g/l FeSO4, 0.005 g/l NaMoO4, and 2 g/l CaCO3) [Bric et al., 1991]. The medium also contained 1 ml of 0.2% L-tryptophan. The cultures were incubated at (25 ± 2) °C for 72 hours with continuous shaking (100 rpm), alongside an uninoculated medium serving as a control. Following incubation, the cultures were centrifuged for 10 minutes at 12,000 rpm, and 1 ml of the clear supernatant was mixed with 2 ml of Salkowsky reagent (comprising 50 ml of 35% perchloric acid and 1 ml of 0.05 mol/L FeCl3 solution). The mixture was then incubated in the dark at room temperature for 30 minutes. The change in color from visible light pink to dark pink indicated IAA production, and the absorbance at 530 nm was measured using a spectrophotometer. The IAA content was calculated using an authentic IAA standard curve.

#### 2.7.2. Screening for inorganic phosphate solubilization by isolated bacteria on agar assay

All bacterial isolates underwent testing for mineral phosphate solubilization activity, employing the National Botanical Research Institute’s phosphate (NBRIP) growth medium supplemented with 1.5% Bacto-agar (Difco Laboratories, Detroit, MI, USA) [42]. Triplicate inoculations of each bacterial isolate were carried out on NBRIP agar medium and incubated for 72 hours at (25 ± 2) °C. The capacity of the bacteria to solubilize insoluble tricalcium phosphate (TCP) was evaluated using the phosphate solubilization index (PSI) [PSI = A / B, where A represents the total diameter (colony + halo zone), and B is the diameter of the colony [43]. The quantification of solubilized phosphorus (P) was determined by subtracting the available P in the inoculated sample from the corresponding uninoculated control [44].

#### 2.7.3. Screening for mineral potash solubilization by isolated bacteria on plate assay

The screening for mineral potassium solubilizing activity in all isolated bacteria was conducted using modified Aleksandrov media [45], incorporating insoluble potassium minerals. Each bacterial isolate was individually inoculated in a petri dish and incubated at 28°C for 7 days post-inoculation. The isolates were cultured in modified Aleksandrov media containing waste biotite at a concentration of 3g/l. The potassium solubilizing bacteria (KSB) were assessed based on the characteristics of their halo zones. After incubation, the measurements of the halo zone and colony diameter were recorded. The potassium solubilization capacity of the isolates was determined using the potassium solubilizing index (KSI) [KSI = A / B, where A represents the total diameter (colony + halo zone), and B is the colony diameter [43]. The quantification of solubilized potassium (K) was calculated by subtracting the available K in the inoculated sample from the corresponding non-inoculated control [44].

#### 2.7.4. Assessment of growth and yield performances of rice grown in nutrient-deficit soil

To assess the plant growth promotion capabilities of the two most effective probiotic bacteria, BTCP01 and BTDR03, a pot experiment was conducted using rice seeds (CV. BRRI dhan29) from November 2016 to May 2017.

The experimental soil, with a slightly acidic pH of 6.41 and clayey texture up to 50 cm depth, contained 0.08% total nitrogen (N), 9 mg/kg available phosphorus (P), 5.7 mg/kg soil-exchangeable potassium (K), and 1.55% organic matter. Meteorological data, including air and soil temperatures at a depth of 30 cm, and rainfall were obtained from the weather archive of the Department of Agricultural Engineering, BSMRAU. Throughout the crop’s growing season, the maximum air temperature ranged from 23.5°C to 36.5°C, while the minimum air temperature ranged from 9°C to 27°C (Fig. S1).

Chemical fertilizers (2.10 g urea, 0.86 g gypsum, and 0.46 g zinc sulfate per 10 kg of soil) were applied based on the Fertilizer Recommendation Guide (FRG) for rice seed CV. BRRI dhan29. Triple superphosphate (TSP) and muriate of potash (MoP) were applied as a basal dose, with urea administered in three equal doses as top dressing at specific growth stages. Cultural practices, including weeding and irrigation, were performed as needed.

Forty-five-day-old seedlings with 3/4 leaves were transplanted into pots (20 cm x 20 cm x 30 cm), with one seedling per pot. The two efficient strains, BTCP01 and BTDR03, were separately grown in 250 ml conical flasks with 200 ml nutrient broth on an orbital shaker for 72 hours. Cells were collected, washed, and prepared as a bacterial suspension. Roots of seedlings were dipped in the bacterial suspension overnight, and freshly harvested bacteria were sprayed on rice plants during the tillering and flowering stages.

The experiment was set in a completely randomized design with three replications which included untreated control i) and treatments with BTCP01 (ii) and BTDR03 (iv) using 0%, 50%, and 100% doses of recommended N, P, and K fertilizers. Essential plant growth parameters were recorded, including root and shoot length, root and shoot dry weight, SPAD value of the flag leaf at panicle initiation, number of tillers and effective tillers per plant, and total grain weight per pot.

### 2.8. Statistical analysis

The data obtained from seedling assays, the P and K solubilizing study and the pot experiments underwent analysis of variance using SPSS (version 17.0) and Statistix (version 10.1). Statistical differences among mean values were determined using the least significant difference (LSD) test at a 5% probability level. The presented data represent mean values ± standard error.

## 3. Results

### 3.1. Isolation, biochemical and molecular characterization of weed endophytic bacteria

A total of 45 bacteria, exhibiting diverse shapes and colors of colonies on nutrient broth agar (NBA) plates, were isolated from surface-sterilized shoots and roots of the collected weeds. These isolates were subsequently purified through repeated streak cultures on NBA medium (Fig. S2). Their impact on seed germination and seedling growth of rice remarkably varied, with some isolates demonstrating inhibitory effects on rice seed germination, as illustrated in Figure S2. Following comprehensive screening, two strains, namely BTCP01 and BTDR03, were chosen based on their superior effects on seed germination rate, shoot length, root length, fresh shoot weight, and fresh root weight of rice (Table S2, Fig. S3, Fig. S4). BTCP01 exhibited a negative Gram reaction, while BTDR03 tested positive (Table 1). Both strains tested positive for catalase and oxidase tests (Table 1).

**Table 1.** Biochemical and molecular characterization of rice probiotic bacteria isolated from different sources.

| Strain | Source of isolation | Biochemical analysis |  |  |  | Molecular analysis |  |  |
| --- | --- | --- | --- | --- | --- | --- | --- | --- |
|  |  | KOH test | Gram reaction | Catalase test | Oxidase test | Accession No. | Closest strain from gene bank | Sequence similarity (%) |
| **BTCP01** | Surface sterilized shoot of Goose grass (*Elusine indica*) | + | - | ++ | + | MW165536 | *Alcaligenes faecalis* | 99% |
| **BTDR03** | Surface sterilized shoot of Bermuda grass (*Cynodon dactylon*) | - | + | ++ | + | MZ798368 | *Metabacillus indicus* | 99% |

‘+’ indicates positive response; ‘-’ indicates negative response.

Phylogenetic analysis based on the constructed tree using 16S rRNA sequences identified the selected strains as members of the genera *Alcaligenes* and *Metabacillus* (Table 1). A BLASTN search at the GenBank database of NCBI revealed that the sequence of BTCP01, deposited under accession number MW165536, exhibited 99% sequence homology with *Alcaligenes faecalis* (Table 2). The sequences of the isolated strain BTDR03, submitted to GenBank under accession numbers MZ798368, displayed 99% similarity with *Metabacillus indicus* (Fig. S5).

**Table 2.**
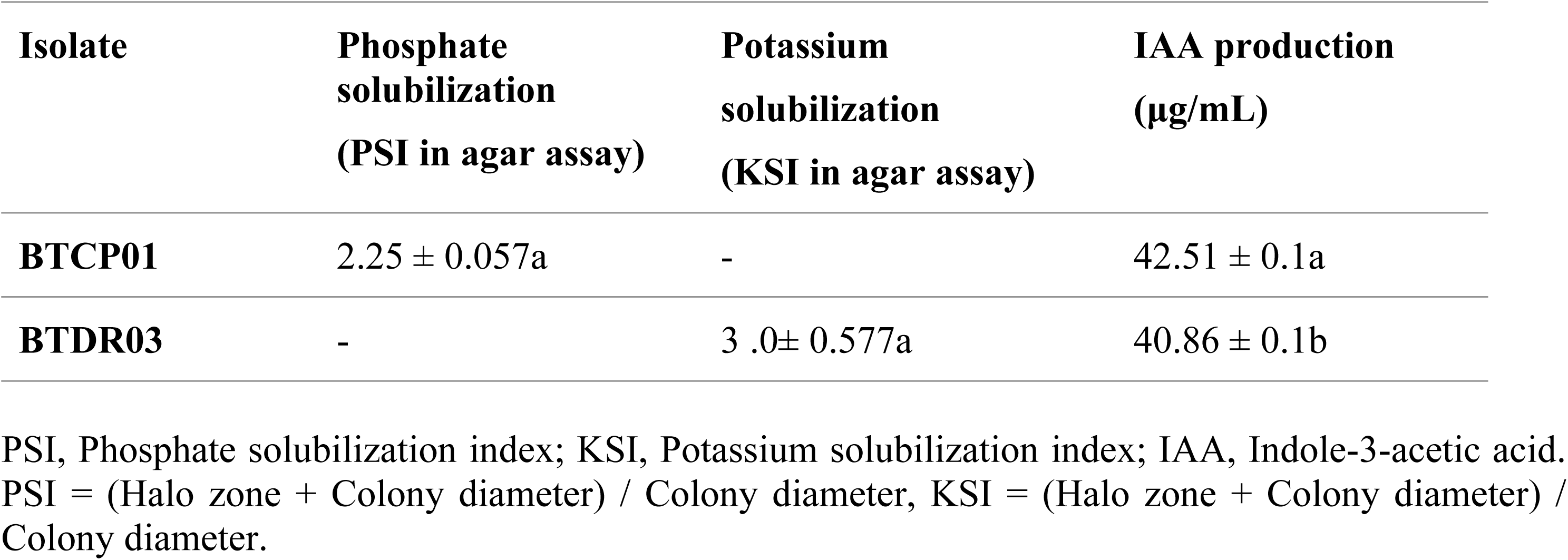
Plant growth-promoting traits of rice probiotic bacteria (Mean ± SE, *n* = 3).

### 3.2. Characterization for plant growth promoting traits of the isolated bacteria

Among the 45 isolates, 20 demonstrated the production of indole-3-acetic acid (IAA) in the presence of L-tryptophan, with concentrations ranging from 13 to 52.78 µg/ml. BTCP01 and BTDR03 displayed IAA production at levels of 42.51 µg/mL and 40.86 µg/mL, respectively (Table 2, Fig. S4C). From this set of isolates, only six exhibited a halo zone on NBRIP agar medium, signifying their phosphate-solubilizing ability. Notably, BTCP01 demonstrated the highest phosphate-solubilizing activity, yielding a PSI value of 2.258 (Table 2, Fig. S4A).

Furthermore, among the 20 isolates, only five displayed a halo zone on modified Aleksandrov media (Hu et al., 2006) containing insoluble potassium minerals. Among these, BTDR03 exhibited the highest potassium-solubilizing index (KSI) with a value of 3.0 (Table 2, Fig. S4-B).

### 3.3. Genetic identity of probiotic bacteria for growth promotion

Both bacterial isolates (BTCP01 and BTDR03), which exhibited varying levels of growth promotion activities such as IAA production, P and K solubilization, and N-fixation, underwent further scrutiny for the presence of key genes regulating these processes. The isolates were subjected to PCR amplification using gene-specific primers (Table S1) to assess the presence of *nifH* (responsible for N-fixation) and *ipdC* (IAA production) genes. Both CPR01 and DRB03 isolates were found to harbor the *nifH* and *ipdC* genes in their genomes (Table 3). These findings suggest that the majority of the isolates produced IAA, partly through the utilization of the indole-3-pyruvic acid (IPyA) pathway.

**Table 3.** Presence (+) or absence (-) of *nifH* and *ipdC*, genes in bacterial genomes.

| Isolates | *ipdC* gene | *nifH* gene |
| --- | --- | --- |
| **BTCP01** | + | + |
| **BTDR03** | + | + |

### 3.4. Promotion of growth and yield of rice cv. BRRI dhan29

The application of BTCP01 and BTDR03 significantly enhanced the growth and yield of rice (Fig. 1 and Fig. 2). The tallest plants were observed when 100% of the recommended chemical fertilizer dose was applied to the plants treated with BTDR03 (116 cm), followed closely by BTCP01 (115.33 cm), surpassing the height of uninoculated plants (109.33 cm) under the same fertilizer dose (Fig. 3A).

**Fig. 1:**
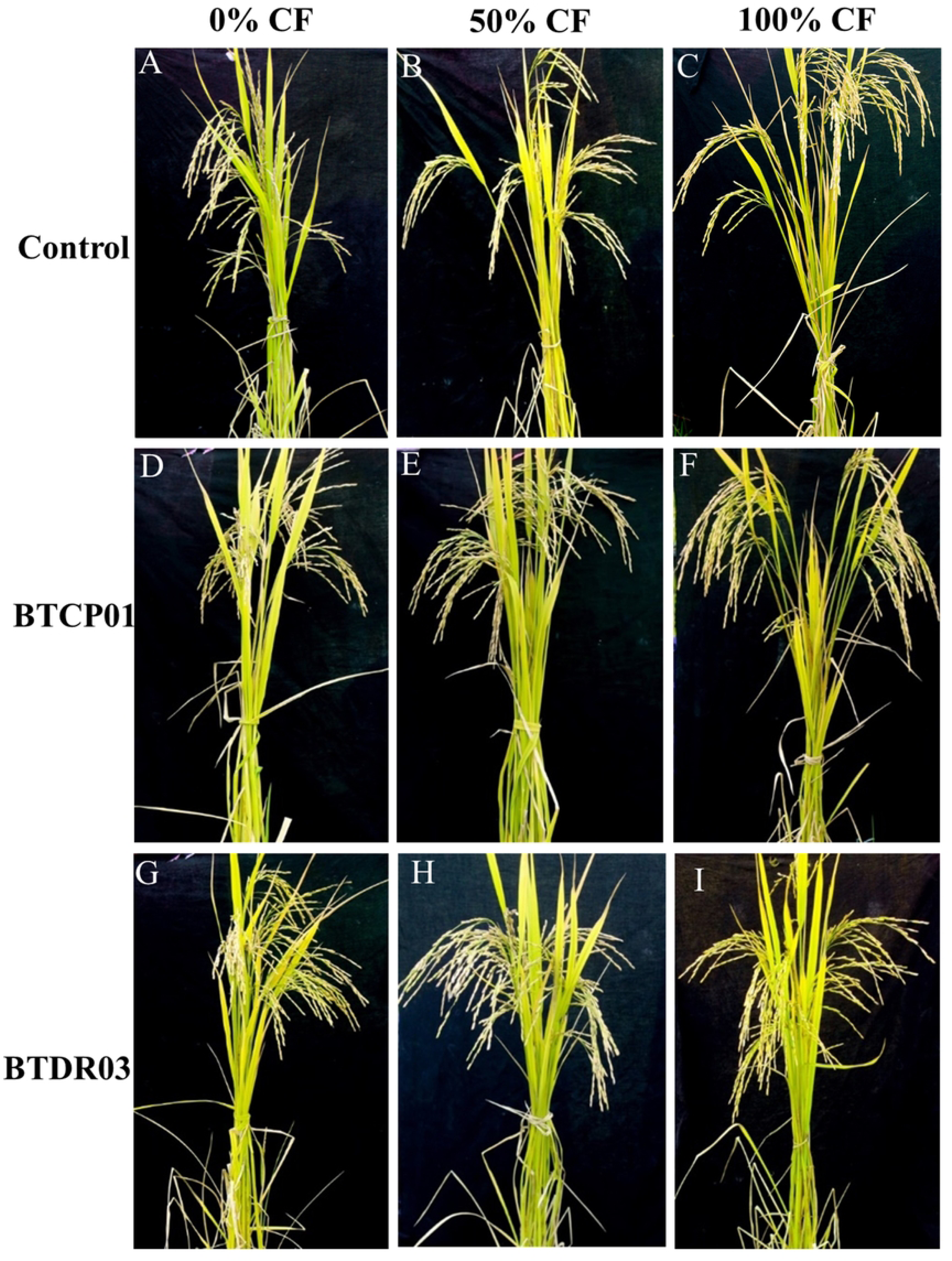
Effects of BTCP01 and BTDR03 on growth performances of CV. BRRI dhan29 with 0% (A, D, C), 50% (D, E, F) and 100% (G, H, I) of the recommended doses of chemical fertilizers respectively. *CF (recommended doses of chemical (N, P, K) fertilizers).

**Fig. 2:**
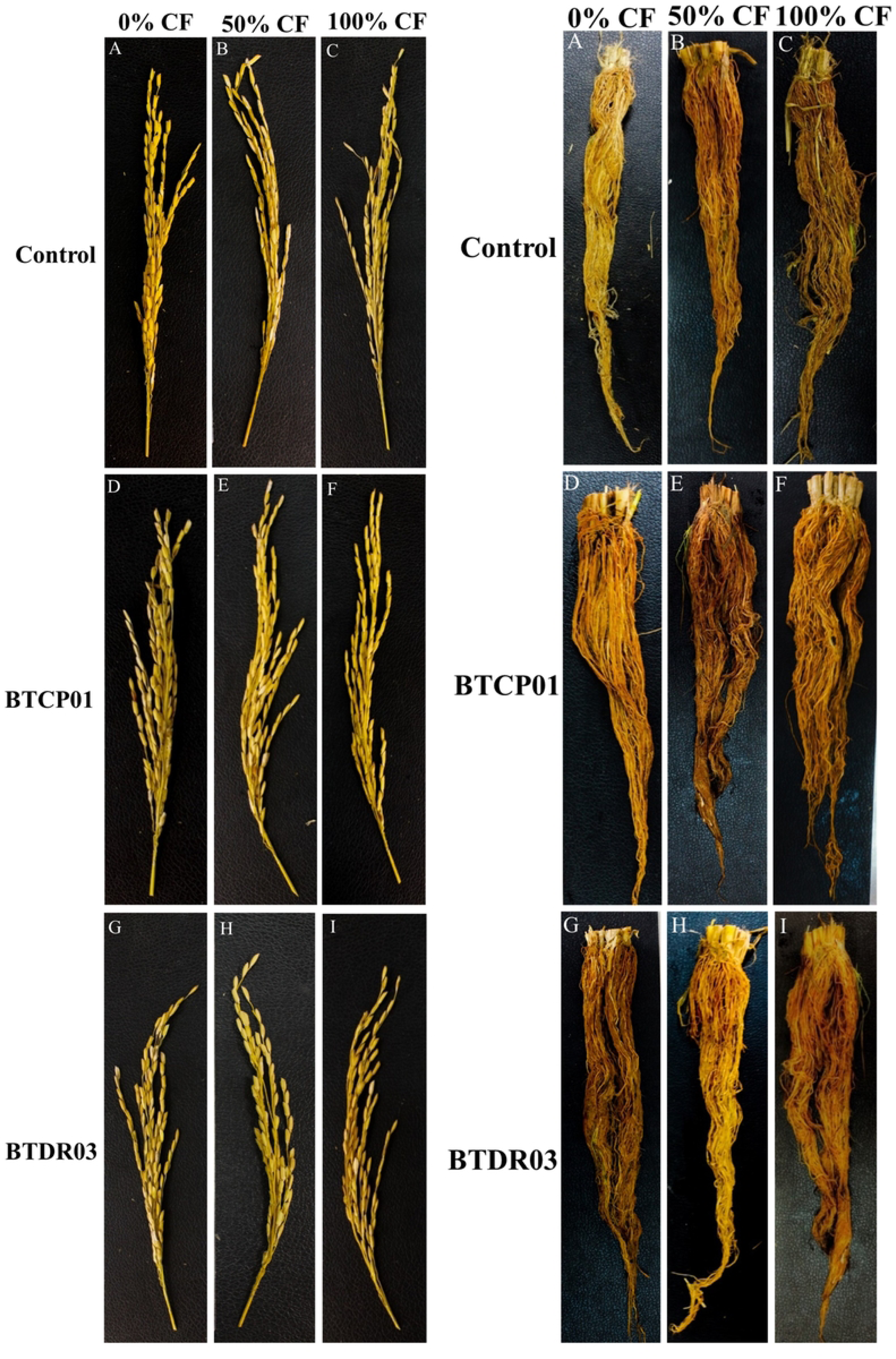
Effects of BTCP01 and BTDR03 on panicle length and root growth performance of CV. BRRI dhan29 with 0% (A, D, C), 50% (D, E, F) and 100% (G, H, I) of the recommended doses of chemical fertilizers respectively. *CF (recommended doses of chemical (N, P, K) fertilizers).

**Fig. 3:**
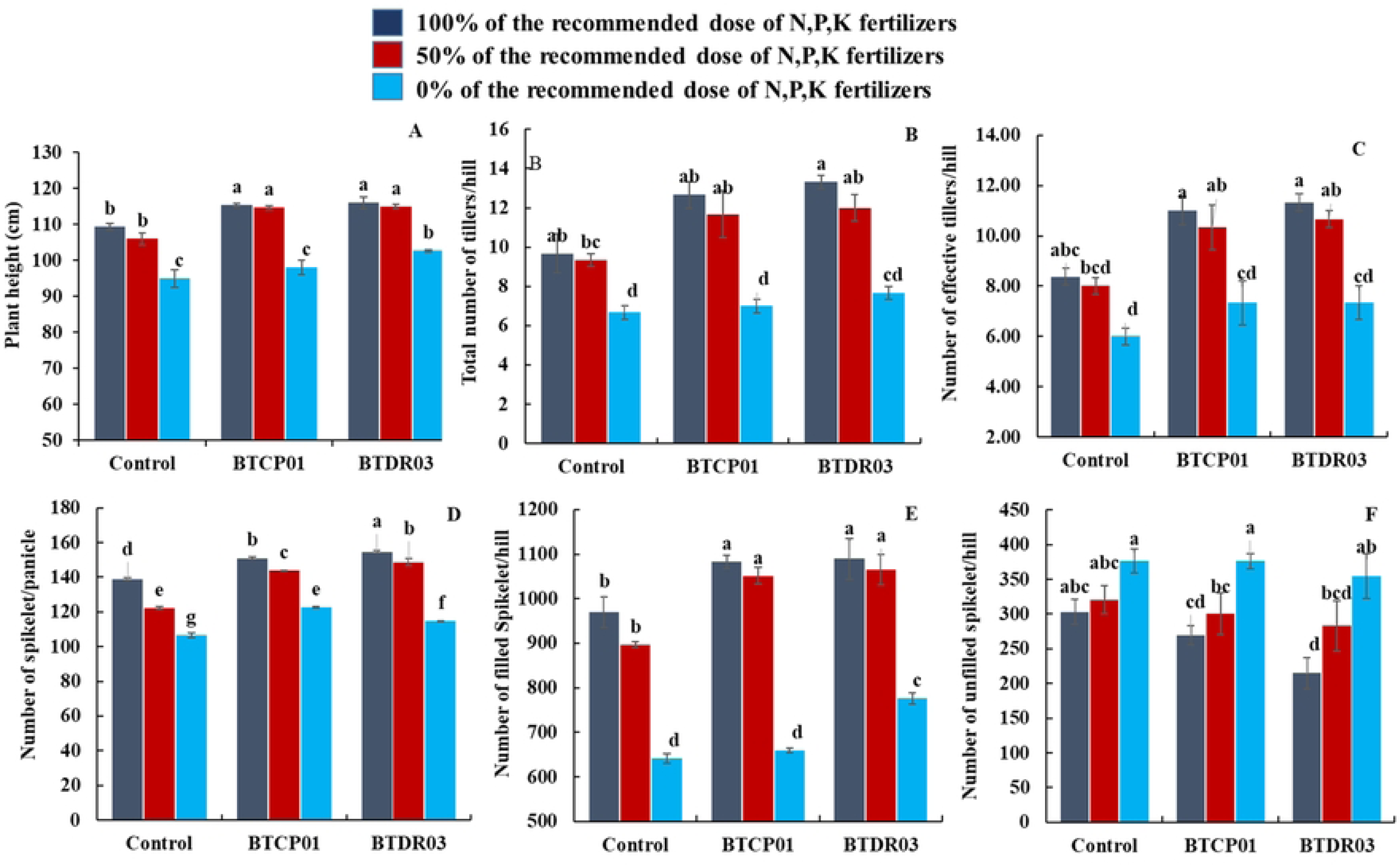
Effects of BTCP01 and BTDR03 along with different fertilizer doses on various growth parameters of BRRI dhan29. (A) Effects of probiotic bacteria on plant height of rice, (B) Effects of probiotic bacteria on total number of tillers per hill of rice, (C) Effects of probiotic bacteria on number of effective tillers per hill of rice, (D) Effects of probiotic bacteria on number of spikelet per panicle of rice, (E) Effects of probiotic bacteria on number of filled spikelet per hill of rice, (F) Effects of probiotic bacteria on number of unfilled spikelet per hill of rice. Values (Mean ± SE, *n* = 3) followed by the same letter(s) in the same graph did not differ significantly at the 0.05 level by the LSD test. Values (Mean ± SE, *n* = 3) followed by the same letter(s) in the same graph did not differ significantly at the 0.05 level by the LSD test.

A noteworthy improvement was observed in various growth parameters, including total tiller number per hill, effective tiller number per hill, number of spikelets per panicle, number of filled spikelets per hill, 1000 grain weight, grain yield (t/ha) per pot, shoot fresh and dry weight, and root fresh and dry weight, in bacterial-inoculated plants (Fig. 3B-F and Fig. 4A-F).

**Fig. 4:**
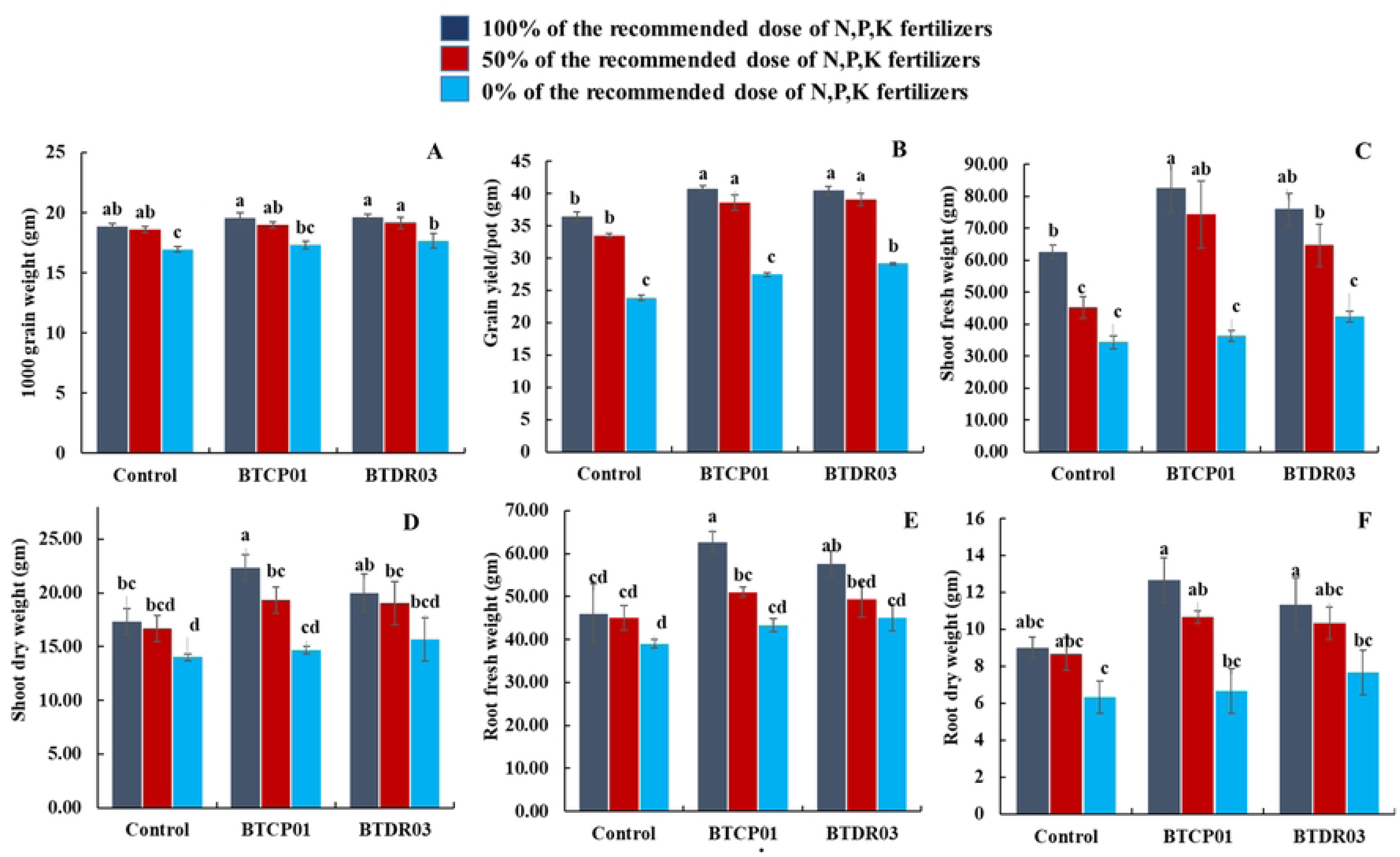
Effects of BTCP01 and BTDR03 along with different fertilizer doses on various growth parameters of CV. BRRI dhan29. (A) Effects of probiotic bacteria on 1000 grain weight (g) of rice, (B) Effects of probiotic bacteria on grain yield per pot of rice, (C) Effects of probiotic bacteria on shoot fresh weight (g) of rice, (D) Effects of probiotic bacteria on shoot dry weight (g) of rice, (E) Effects of probiotic bacteria on root fresh weight (g) of rice, (F) Effects of probiotic bacteria on root dry weight of rice. Values (Mean ± SE, *n* = 3) followed by the same letter(s) in the same graph did not differ significantly at the 0.05 level by the DMRT test using LSD parameter.

Application of 100% of the recommended chemical fertilizer dose to BTCP01 and BTDR03-treated plants significantly increased total tillers per hill and effective tillers per hill compared to untreated controls receiving an equal fertilizer dose (Fig. 3C, D). Notably, the treatment with 50% of the recommended fertilizer dose along with BTCP01 resulted in more than a 20.68% increase in total tillers per hill (11.66) and a 23.49% increase in effective tillers per hill (10.33). Similar increases were observed for BTDR03, with more than 24% in total tillers per plant (12) and around 27% in effective tillers per plant (10.667) compared to uninoculated plants receiving 100% of the recommended chemical fertilizer dose (Fig. 3B, C).

The highest number of spikelets per panicle was achieved when 100% of the recommended chemical fertilizer dose was applied to BTDR03-treated plants, followed by BTCP01, with both slightly surpassing the uninoculated control grown with the same fertilizer dose (Fig. 3D). Applying 100% of the recommended chemical fertilizer dose to BTDR03-treated plants produced the highest number of filled spikelets (1089), followed by BTCP01 (1082), significantly surpassing the uninoculated control (970) grown with the same fertilizer dose (Fig. 3E). Notably, around 8.31% and 9.78% increases in filled spikelets were observed in BTCP01 and BTDR03-treated plants with 50% of the recommended fertilizer dose, respectively, compared to the untreated control receiving 100% of the recommended fertilizer dose (Fig. 3E). Moreover, BTCP01 and BTDR03-treated plants with 50% of the recommended fertilizer dose exhibited a notable 5.64% and 6.75% increase in rice grain yield per pot, respectively, compared to the control treatment (Fig. 4B). A similar increasing trend, although not statistically significant, was observed in BTCP01-treated plants, while a significant increase was noted in BTDR03-treated plants with 0% of the recommended fertilizer dose, suggesting the potential to reduce major fertilizer use in rice production by up to 50% (Fig. 4B).

In addition to grain yield, shoot fresh and dry weight of rice substantially increased in BTCP01-treated plants under 100% of the recommended fertilizer dose, surpassing the untreated control grown under the same conditions (Fig. 4C, D). Similar trends were observed in treatments with BTCP01 using 100% of the recommended fertilizer dose, significantly enhancing root fresh and dry weight compared to uninoculated control plants grown with similar fertilizer doses (Fig. 4E, F).

## 4. Discussion

In this study, we isolated 45 endophytic bacteria from native rice weeds and identified two promising rice growth-promoting bacteria, *Alcaligenes faecalis* (BTCP01) and *Metabacillus indicus* (BTDR03), through 16S rRNA gene sequencing. These diazotrophic bacteria were found to significantly enhance seed germination, seedling growth, and ultimately the yield of rice, even with a 50% reduction in N, P, and K fertilizers. We established that the growth-promoting effects were associated with nitrogen fixation (N-fixation), indole-3-acetic acid (IAA) production, and the solubilization of mineral phosphates and potash, suggesting key elements contributing to plant growth and productivity [46]. While numerous growth-promoting endophytic bacteria have been previously isolated from various plant sources [12,13,46], few have displayed harmful impacts on seed germination and growth. This study identified weed endophytes that significantly improve rice growth and yield, showcasing their potential to reduce fertilizer use by up to 50%, without compromising yield.

A notable discovery in our study was the isolation of diazotrophic *A. faecalis* bacterium (BTCP01) from Goose grass (*Eleusine indica*), a native rice weed, which remarkably increased rice growth and yield with a 50% reduction in major chemical fertilizers (Table 1 and Figs. 1-4). *A. faecalis* (BTCP01), initially discovered in feces, has been isolated from various environments, demonstrating its potential as a plant growth-promoting bacteria (PGPB) [48-54]. Our findings indicated that BTCP01 promoted rice growth and yield through nitrogen fixation (detected by *nifH* gene) and IAA production (detected by *ipdC* gene). *A. faecalis* has previously been reported for phosphorus solubilization in strains isolated from *Nicotinia glutata*, further supporting its potential as a multifaceted PGPB [57]. It has also demonstrated efficacy as a halotolerant PGPB, aiding the vegetative development of salinity-stressed rice, wheat, and canola plants [58-60].

One of the interesting findings of our study was that the application of BTCP01 and BTDR03 with reduced fertilizer doses resulted in statistically equal or higher root length, total tillers per hill, effective tillers per hill, and grain yield compared to untreated control plants receiving 100% of the recommended doses (Figs. 3 and 4). These findings suggest that these two weed endophytic bacteria could effectively reduce N-P-K fertilizer use by up to 50% without compromising rice growth and yield. Coinoculation with various strains of *Burkholderia* spp. and *Pseudomonas aeruginosa* from different weeds has been reported to enhance plant growth and yield in several crops, emphasizing the potential for field evaluations [65-68]. However, we for the first time demonstrated that weed endophytic bacteria *A. faecalis* and *M. indicus* isolated from the rice weeds have potential for reduction in chemical fertilizers in rice. A further field level evaluation of these two bacteria either alone or in combination are needed to confirm the potentials as candidates for biofertilization in rice.

To see the mechanistic insights of the higher yield in rice by the weed endophytic bacteria, we checked whether they possess any genetic traits in their genome association with plant growth promotion. Interestingly, we detected *nifH* and *ipdC* genes in the genomes of BTCP01 and BTDR03 that suggests their potential to fix nitrogen and produce phytohormone IAA. While the growth and grain yield enhancement of treated plants may be associated with increased nutrient uptake [69], additional mechanistic studies are needed to uncover the full potential of these bacterial isolates as bioinoculants. Notably, some weed endophytes in our study severely suppressed rice seed germination (Table S2). Given that weeds are known competitors of rice and can inhibit seed germination and crop growth, understanding the molecular basis of weed endophytes inhibiting rice seed germination warrants further investigation. Although allelopathic effects of weeds through phytotoxic secondary metabolites have been reported [70], reports on allelopathic effects of weed endophytes are limited. Phytotoxic compounds from weed endophytes may offer new herbicide candidates. Our findings underscore the importance of discovering novel weed endophytes as valuable bioresources for sustainable agriculture, contributing to a reduction in the use of synthetic agrochemicals that pose threats to soil and environmental health [7].

In conclusion, this study represents the first identification and characterization of two weed endophytic bacteria, *A. faecalis* and *M. indicus,* with the ability to enhance the growth and yield of rice under 50% reduced doses of N, P, and K chemical fertilizers. The underlying mechanisms of their beneficial effects encompass IAA production, atmospheric N-fixation, and mineral P and K solubilization. Additionally, the presence of *nifH* and *ipdC* genes correlates with their growth-promoting activities on rice. Consequently, these endophytic bacteria, sourced from rice-associated weeds, hold significant promise for reducing chemical fertilizer usage in sustainable rice production. However, a comprehensive multi-location field evaluation of these strains is imperative before recommending them as biofertilizers for rice cultivation. Moreover, exploring the synergistic effects of coinoculating these two weed endophytes on rice presents an intriguing avenue for further investigation.

## Used abbreviations

IAA: Indole-3-acetic acid
PGPB: plant growth-promoting bacteria
NB: Nutrient broth
KSB: potassium solubilizing bacteria
PSB: phosphate solubilizing bacteria
PSI: phosphate solubilizing index
KSI: potassium solubilizing index.

## Data availability statement

The original contributions presented in the study are included in the article/supplementary materials, further inquiries can be directed to the corresponding author.

## Author contributions

**Kaniz Fatema; Nur Uddin Mahmud:** Conceptualization, Methodology, Formal analysis, Investigation, Data curation, Writing - original draft, Writing - review & editing. **Dipali Rani Gupta; Md Nurealam Siddiqui; Tahsin Islam Sakif; Aniruddha Sarker; Andrew G Sharpe:** Writing - original draft, Writing - review & editing. **Tofazzal Islam:** Conceptualization, Methodology, Writing - review & editing, Supervision, Funding acquisition.

## Acknowledgement

The authors are thankful to Bangladesh Academy of Sciences (BAS) for the funding of this work under a project No. BAS-USDA PALS CR-11.

## Supplementary material

The supplementary material for this article can be found online.

